# Clonal analysis reveals differential PGC contributions to the early germline and ovarian reserve

**DOI:** 10.1101/2025.04.30.651497

**Authors:** Maya Pahima, Florence L. Marlow

## Abstract

Primordial germ cells (PGCs) are the precursors of the germline and among the first cells to be specified during embryogenesis. Contrary to the common assumption that PGCs directly develop into germline stem cells (GSCs), recent mammalian data and our zebrafish observations suggest a more complex picture. We hypothesized that individual PGCs contribute differentially to gonad development. To test this hypothesis, we developed two Cre-based lineage tracing systems and demonstrate that PGCs give rise to at least four distinct clone types in zebrafish. We observed no restriction in clone type localization along the anterior-posterior axis, suggesting these differences are PGC intrinsic or locally determined. Further examination of the ovarian reserve revealed functional evidence that some PGCs generate self-renewing GSCs, while others produce non-renewing progenitors that are depleted with successive matings. Collectively, our findings propose a revised model of vertebrate germline establishment with implications to reproductive lifespan. This model suggests that differentiation potential varies among PGCs, allowing individual PGCs to directly differentiate into multiple distinct germ cell types, including renewing GSCs and transient populations that contribute to an early wave of gametes.

## Introduction

In zebrafish, the primordial germ cells (PGCs) are specified by inheritance of maternal germ plasm — aggregates of germline-specific mRNAs and proteins allocated during the first cell divisions (reviewed in (Marlow and 2015)). 4 cells initially inherit the germ plasm and, through subsequent divisions, establish a founding population of ∼25-50 PGCs (Braat et al., 1999; Knaut et al., 2000; Raz, 2002; Weidinger et al., 1999; Yoon et al., 1997). PGCs are distinctly larger than surrounding cells and can be identified by the highly conserved molecular marker Ddx4 (Vasa) (Braat et al., 1999; Knaut et al., 2000; Weidinger et al., 1999; Yoon et al., 1997). PGCs migrate to the gonadal ridge in the first 30 hours post-fertilization (30h) (Raz, 2002; Raz et al., 2006), where they remain until differentiation at day 7 (d7) (Bertho et al., 2021) (Fig. 1A). By d14, the PGCs have undergone differentiation to form the early germline, also known as the pre-gametic germline (Ogielska et al., 2024). The pre-gametic germline is comprised of smaller, uniformly-sized germ cell daughters and a few germline stem cells (GSCs) identifiable by expression of *nanos2* (Bertho et al., 2021). Around d19, some germ cells undergo meiotic entry and begin to develop into nascent oocytes (Dranow et al., 2016). At this stage, the gonad is considered “bipotential” and expresses genes involved in both testis and ovary development (Kossack and Draper, 2019; Leerberg et al., 2017; Rodriguez-Mari et al., 2005). Upon initiation of sex determination and changes in the somatic gonad fates between d25-d28, the gonad will finally develop into either an ovary or testis (Dranow et al., 2016; Rodríguez-Marí et al., 2010).

**Fig. 1:**
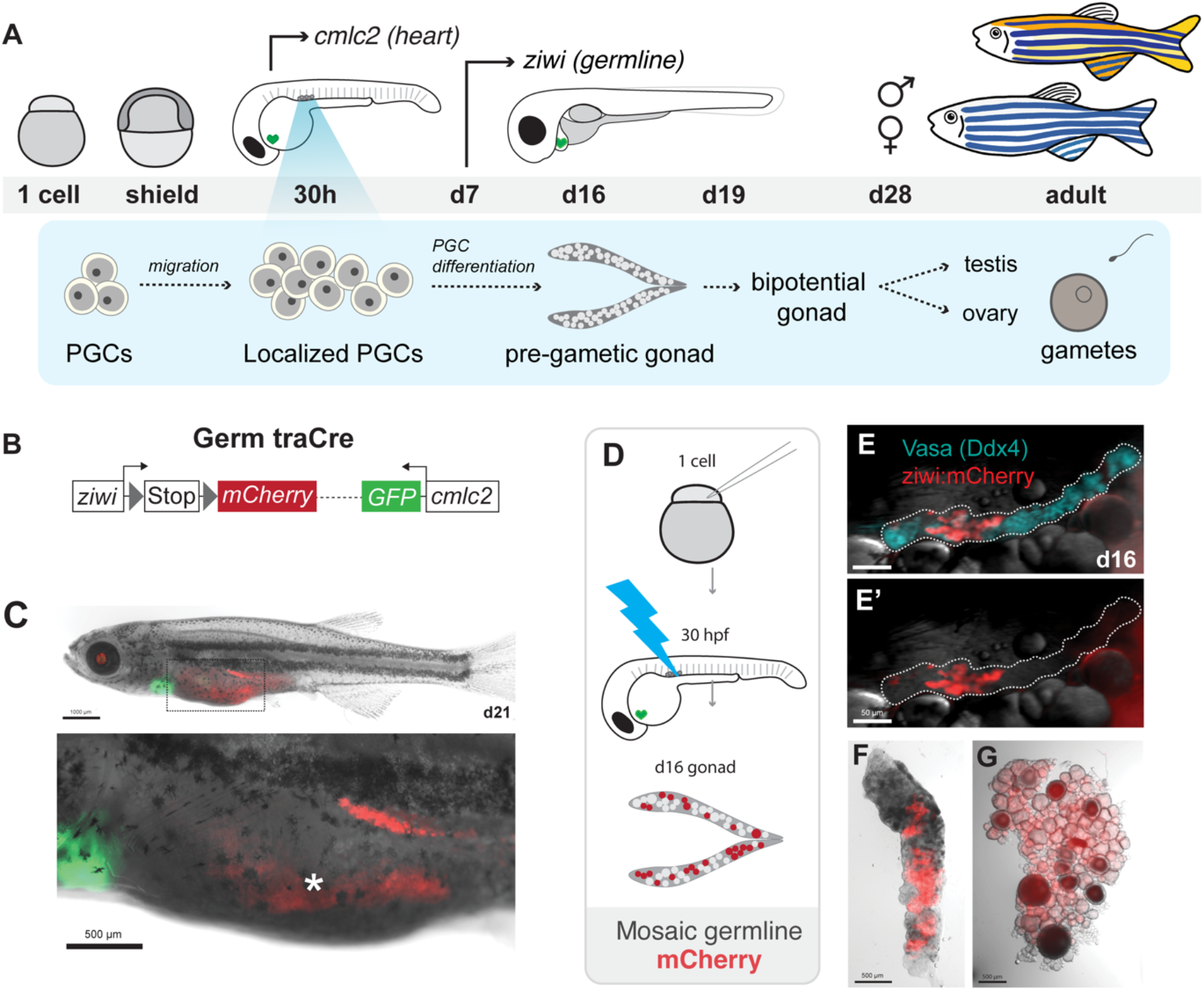
Zebrafish germline development and a novel germline lineage tracing system, Germ traCre. **(A)** A schematic overview depicting zebrafish germline development, from PGCs to mature gametes. Arrows indicate developmental timing of the *ziwi* and *cmlc2* promoters used in transgenic constructs. (**B)** Schematic of the Germ traCre transgenic constructs and photoactivatable Cre (pACre) with *nos3, 3’utr* control element for germline-specific translation. **(C-C”)** Germ traCre transgenic lines were validated by injection of Cre mRNA. (C) Juveniles were examined at d21 for broad germline-specific mCherry expression (red) and pGH green heart marker expression. Scale bar: 1000 μm. Dashed box outlining enlarged image of heart and gonad expression. **(C’-C”)** Enlarged heart and germline view. Gut autofluorescence indicated by asterisk. Scale bar: 500 μm. **(D)** Schematic of the Germ traCre experimental workflow used to generate mosaic germline labeling. pACre-*nos3,3’utr* mRNA was injected into one-cell embryos and activated with blue light at 30h to induce recombination and marker expression. **(E-G)** Mosaic Germ traCre gonads. (**E-E’**) d16 gonadal lobe with bright field imaging, mosaic mCherry expression (red), and Ddx4 germline labeling (blue). Scale bar: 50 μm. **(F-G)** Mosaic mCherry expression in adult ovary and testis (∼100d). Scale bars: 500 μm.

### PGC differentiation

In many animals, the PGCs establish the germline prior to sexual determination. Thus, early germ cells must have the potential to support development of either the male or female germline. Given this, PGCs have largely been assumed to be germline stem cells (GSCs) or to give rise to sexually-indifferent GSCs which then differentiate to establish a bipotential juvenile gonad (reviewed in (Tanaka, 2016)). However, despite their common origin from a sexually indifferent gonad, mammalian testes harbor a population of GSCs that ovaries lack (Saba et al., 2014; Sada et al., 2009) and recent sequencing data shows that zebrafish PGCs and GSCs are distinct cell states with heterogeneous populations (Gross-Thebing et al., 2017; Zhang et al., 2019). Moreover, we recently showed that PGCs are more numerous than the initial number of GSCs in the early zebrafish germline (Bertho et al., 2021) — from ∼25 PGCs on one side of the gonad, only 1-2 of their daughter cells express the conserved GSC marker *nanos2* (*nos2)* (Cao et al., 2019; Sada et al., 2009). These observations led us to hypothesize that not all PGCs give rise to GSCs and may instead differentially contribute to the early germline.

Investigating vertebrate PGC contributions to the germline is challenging. Previous studies in zebrafish have labeled PGCs using the germ plasm mRNA UTR elements that restrict mRNA stability and translation to PGCs (Köprunner et al., 2001; Saito et al., 2006; Suzuki et al., 2010). Although leveraging the maternal mechanism of germ plasm specification is an elegant approach to label the PGCs, it provides only a transitory view of their development. This is because PGCs are both the earliest cell type specified in the embryo, and some of the latest embryonic cells to differentiate, beginning at d7 (Bertho et al., 2021). Thus, the maternally-inherited factors and injected mRNAs traditionally used to label PGCs cannot be reliably used to lineage trace individual PGC germline contributions. To circumvent this problem, we established methods to lineage trace zebrafish PGCs and their progeny.

In this study, we developed Germ traCre and Germbow — two Cre-based transgenic lineage tracing systems to investigate PGC germline contributions. By combining these genetic marker approaches with hybridization chain reaction fluorescence *in situ* hybridization (HCR FISH) and immunohistochemistry (IHC) labeling of known germ cell markers, we uncovered evidence that individual PGCs can directly differentiate into multiple distinct germ cell types. Further examination of PGC contributions over reproductive lifetime revealed that some PGCs give rise to renewing GSCs and others to non-renewing progenitors which are depleted in the first rounds of mating. Altogether, we propose a revised view of PGC contributions to the early gonad and the implications over reproductive lifespan.

## Results

### Germ traCre: a transgenic system for clonal germline lineage tracing

To examine PGC contributions to the early gonad and GSC pool, we developed Germ traCre: a Cre-based transgenic lineage tracing system for clonal analysis of PGC derivatives, *Tg[ziwi:stop-mCherry; cmlc2:GFP]* (Fig. 1B). We used the zebrafish germline-specific promoter *ziwi*, which is active in germ cells beginning at d7 (Leu et al., 2010; Tan et al., 2002) (Fig. 1A), to drive an mCherry coding sequence silenced by a floxed stop. Germ traCre also contains an independent transgenesis reporter expressed under a cardiac-specific promoter (*cmcl2:GFP)*, enabling larval screening after 30h. As expected, we found that ubiquitous Cre expression induced recombination and stable mCherry expression specifically in germ cells (Fig. 1C).

To label PGCs and produce mosaic gonads with distinct, spatially resolved clonal clusters, we generated a blue light photoactivatable Cre recombinase (pACre) (Kawano et al., 2016) fused to the *nanos3,3’utr* (Köprunner et al., 2001; Suzuki et al., 2010) to restrict mRNA translation to the germline (Fig. 1B). We reasoned that injection of zygotes with *pACre-nos3,3’utr* mRNA would provide efficient spatiotemporal control of Cre activation in germ cells during the period between maternal PGC specification and zygotic *ziwi* activation (∼d7). To generate stable mosaic mCherry+ gonads for clonal analysis of individual PGC contributions over reproductive development, we injected Germ traCre zygotes with a low concentration of *pACre-nos3,3’utr* mRNA and activated the recombinase by exposing embryos at 30h to 470nm blue light (Fig. 1D). Co-expression of mCherry with the germ cell-specific protein Ddx4 confirmed specific labeling of the germline at d16 and in adult ovary and testis. (Fig. 1E). We then combined Germ traCre with HCR FISH and IHC staining for known germ cell markers to examine individual PGC fates.

### PGCs give rise to multiple distinct germ cell fates

To determine if all PGCs differentiate as GSCs, we injected Germ traCre transgenic zygotes with *pACre-nos3,3’utr* mRNA then exposed dechorionated 30h embryos to 470nm blue light for 3 mins to activate the Cre and produce mosaic germline labeling. We screened gonads just after PGC differentiation at d16 to capture small clones (2-4 cells) and identified 99 spatially isolated mCherry+ clones across 15 gonads (Fig. 2A).

**Fig. 2:**
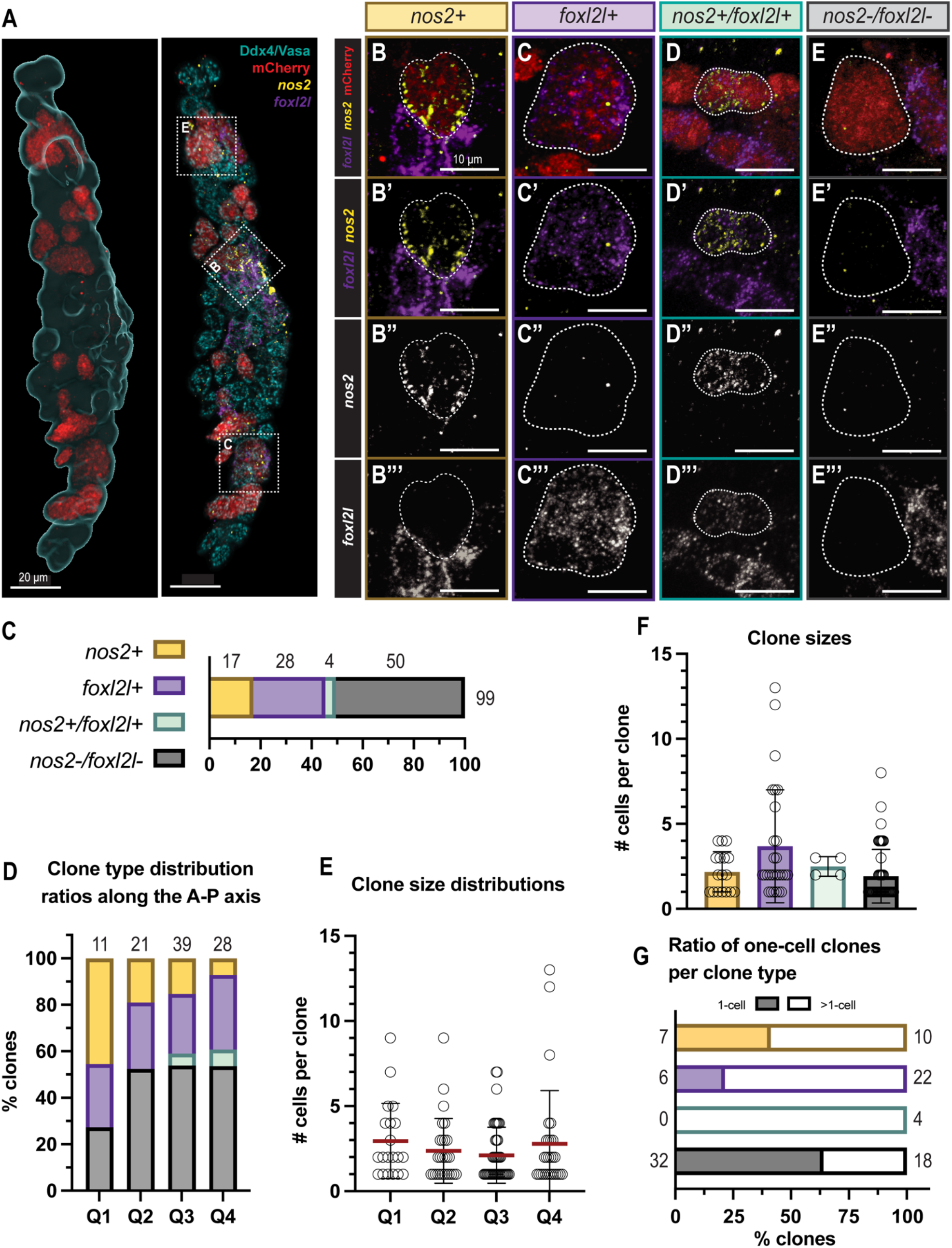
Germ traCre clonal analysis reveals direct descendants of PGCs. **(A-A’)** A d16 mosaic Germ traCre gonadal lobe with mCherry+ clones, with additional IHC and HCR labeling of Ddx4 IHC labeling (Vasa; germline, blue), *nos2* (GSC, yellow), and *foxl2l* (progenitor, purple). **(A)** Imaris software analysis representation with volume rendering of the germline (blue) and mCherry+ clones (red) within. **(A’)** Example *nos2*+, *foxl2l*+, and *nos2-/foxl2l-* clones (B,C,E) marked with dashed boxes. Scale bars: 20 μm. **(B-E”‘)**. Representative images of the four identified clone types. mCherry+ clones are outlined in white dashed lines. Combined and individual channels shown for mCherry, *nos2*, and *foxl2l*. Scale bars: 10 μm. **(C)** Bar graph showing clone type ratios of 99 analyzed clones from 15 co-stained gonads (*nos2*+: 17, *foxl2l*+: 28, *nos2*+/*foxl2l*+: 4, *nos2*-/*foxl2l*-: 50). **(D)** Bar graph showing the distribution ratios of clone types along the gonadal AP axis. To quantify this, gonads separated into 4 quadrants (Q1-Q4) for analysis. Total number of clones per quadrant labeled. **(E)** Scatter plot of clone size distributions along the gonadal AP axis. Means indicated in red with black SD bars (Q1: 2.95 ± 2.21, Q2: 2.37 ± 1.90, Q3: 2.11 ± 1.65, Q4: 2.79 ± 3.13). **(F)** Scatter bar plot of clone sizes between clone types, SD bars indicated (*nos2*+: 17, 2.18 ± 1.19; *foxl2l*+: 28, 3.68 ± 3.23; *nos2*+/*foxl2l*+: 4, 2.50 ± 0.58; *nos2*-/*foxl2l*-: 50, 1.92 ± 1.58). **(G)** Bar plot showing ratio of one-cell clones to multi-cell clones per clone type (*nos2*+: 7:10, 41.18%; *foxl2l*+: 6:22, 21.43%; *nos2*+/*foxl2l*+: 0:4, 0%; *nos2*-/*foxl2l*-: 32:18, 64%).

Given that zebrafish PGCs are thought to directly produce GSCs which later differentiate into a bipotential gonad with nascent oocytes, we expected early gonads to contain heterogeneous clones comprised of individual GSCs and their immediate differentiated daughters. To distinguish GSCs from their more differentiated progeny, we co-labeled mosaic Germ traCre gonads with the highly conserved GSC marker *nos2* (Cao et al., 2019; Sada et al., 2009) and the early differentiation marker *foxl2l* (previously *foxl3*) (Hsu et al., 2024; Liu et al., 2022; Ren et al., 2024) (Fig. 2A). We also used the germ cell marker Ddx4 (Vasa) (Raz and 2000) to better delineate individual cells. To our surprise, none of the examined clones contained a mixture of *nos2*+ and *foxl2l*+ cells. Instead, our clonal analysis revealed homogenous clones with 4 distinct expression profiles: (1) *nos2*+, (2) *foxl2l*+, (3) *nos2+* and *foxl2l+* (*nos2*+/*foxl2l*+), and (4) clones which expressed neither marker (*nos2*-/*foxl2l*-) (Fig. 2B). Of 99 mCherry+ clones analyzed across 15 gonads, we identified 17 *nos2*+, 28 *foxl2l*+, 4 *nos2+/foxl2l+*, and 50 *nos2-/foxl2l-* clones (Fig. 2C). Consistent with our previous observation that the number of *nos2*+ cells in the pre-gametic gonad is fewer than expected based on the initial number of PGCs (Bertho et al., 2021), less than 20% of clones were *nos2+*. Thus, in contrast with previous assumptions about vertebrate PGC differentiation, it appears that the majority of PGCs give rise to clones of non-GSC germ cells without transitioning through a GSC intermediate.

### GSCs form throughout the pre-gametic gonad

In invertebrates such as *Drosophila* and *C. elegans*, PGC fates are influenced by spatially-restricted somatic gonadal niches which induce and maintain the GSC fate (Crittenden et al., 2006; Gilboa and Lehmann, 2004; Sheng et al., 2009). In these organisms, the PGCs that localize near the somatic GSC niche at the anterior end of the gonad differentiate into GSCs, whereas more distantly located PGCs instead differentiate into gonocyte progenitors. To test whether PGC fates in the early zebrafish gonad might be similarly determined by a spatially-restricted somatic GSC niche along the anterior-posterior (AP) axis, we analyzed the spatial distribution of clones by subdividing the d16 gonads into 4 quadrants along the AP axis and quantifying the clone types (Fig. 2D) and clone sizes (Fig. 2E) within each quadrant. We found that all clone types were present in all quadrants along the AP axis (Fig. 2D) with no appreciable difference in clone size distribution (Fig. 2E). These data suggest that zebrafish GSC fate acquisition is not determined or restricted by a cell’s position along the gonadal AP axis.

The presence of GSCs along the AP axis is consistent with zebrafish adult ovary organization. Unlike *Drosophila* and *C. elegans* ovaries, the adult zebrafish ovary has a mitotic germinal ridge within a lateral band that spans AP axis along the surface of the gonad (Beer et al., 2013). Prior wholemount analysis of d21 gonads labeled for *nos2* indicated that GSCs are distributed throughout the gonad (Beer and Draper, 2013). Similarly, our quantification of *nos2*+ cluster number and position across 23 d16 gonads revealed an average of 2.45 *nos2*+ clusters that were present throughout the gonad, although most commonly at the periphery (Fig. S1). Thus, we conclude that the capacity to induce or adopt GSC fate is not restricted along the gonad axes in zebrafish. It remains to be determined whether zebrafish GSC fate specification requires interactions with the somatic gonad.

### The pre-gametic gonad is comprised of multiple progenitor populations

We next compared clone sizes between clone types (Fig. 2F). *nos2*+ clones at this stage had an average of 2 cells, with the largest clone containing 4 cells. Overall, *foxl2l*+ and *nos2-/foxl2l-* clones were the largest: containing as many as 13 and 8 cells, respectively. Yet, while *foxl2l*+ clones had an average size of 3.68 cells, more than half of *nos2-/foxl2l-* clones were unicellular, reducing the average size of this clone type to 1.92 cells. It is possible that unicellular *nos2-/foxl2l-* clones represent undifferentiated PGCs; however, 18/50 (or 36%) of *nos2-/foxl2l-* clones had two or more cells (Fig. 2G), indicating that at least some *nos2-/foxl2l-* clones represent differentiated PGC progeny.

To determine if the *nos2-/foxl2l-* clones were more differentiated germ cells that had entered meiosis and stopped expressing *foxl2l*, we examined the expression of *nos2, foxl2l*, and the meiotic entry factor *rec8a* in WT gonads (Fig. 3A). Although we found some *foxl2l*+ germ cells that co-expressed *rec8a*, no *nos2-/foxl2l-* germ cells had detectable *rec8a* expression at this stage (Fig. 3A). This indicates that *nos2-/foxl2l-* clusters are not meiotic cells that no longer express *foxl2l*. Thus, we conclude that *nos2-/foxl2l-* expressing clones represent an unexpected germ cell fate. Notably, *nos2-/foxl2l-* clones accounted for half of all examined clones (Fig. 2A).

**Fig 3.**
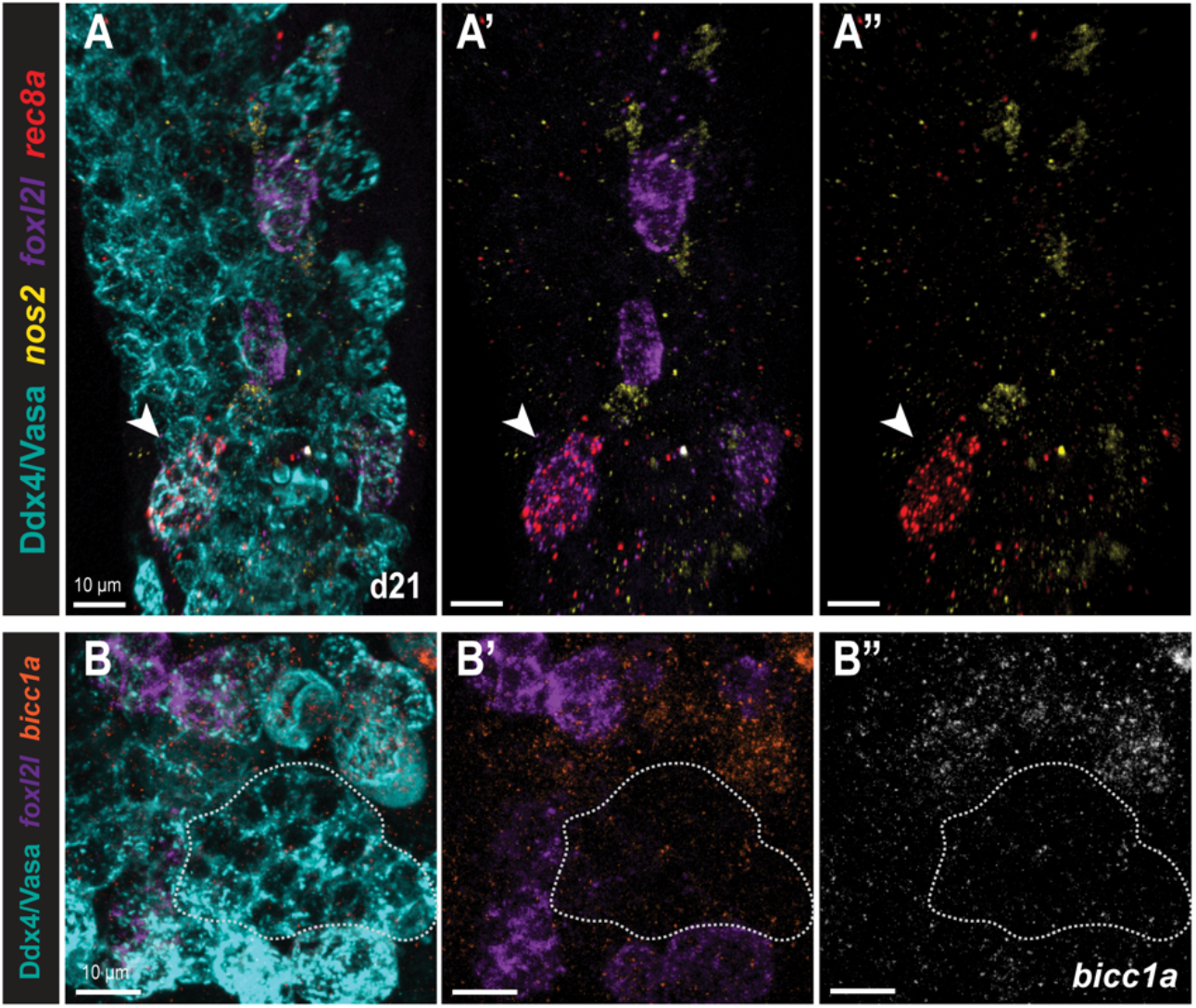
*nos2-/foxl2l-* cells are not meiotic. **(A-A”)** d21 WT gonad stained for Ddx4 (Vasa; germline, blue), *nos2* (GSC, yellow), *foxl2l* (progenitor, purple), and *rec8a* (meiotic entry, red). White arrow: cluster co-expressing *foxl2l*+/*rec8a*+. (n= 3 gonads examined). **(B-B”)** d28 WT gonad stained for Ddx4 (Vasa; germline, blue), *foxl2l* (progenitor, purple), and *bicc1a* (orange). White outline: cluster of *foxl2l*-cells of similar size to *foxl2l*+ cells. (n= 3 gonads examined). Scale bars: 10 μm.

Recent single cell RNA sequencing (scRNAseq) of the d26 bipotential gonad identified distinct progenitor types, including a population of *nos2-/foxl2l-* progenitors similar to our clones (Hsu et al., 2024). Based on the UMAP trajectories, *nos2-/foxl2l-* cells were proposed to arise from GSCs and represent an intermediate state between GSCs and *foxl2l+* progenitors. In the available scRNAseq data, the *nos2-/foxl2l-* cell cluster (referred to as the “Progenitor E” population) appear to express the RNA-binding protein *bicc1a* (Hsu et al., 2024). To determine if our *nos2-/foxl2l-* progenitor clusters were “E-type” progenitors, we examined the expression of *foxl2l* and *bicc1a* in d28 WT gonads. Although we did not detect distinct *foxl2l*+ and *bicc1a+* cell clusters, *bicc1a* mRNA appeared to be present at low levels throughout the germline and was enriched in early oocytes (Fig. 3B). Thus, *bicc1a* does not appear to mark a distinct population of early germ cells at this stage. The identity of the *nos2-/foxl2l-* cells, how many germ cell types they represent, and their fate and renewal capacity remain to be determined.

### Germbow: a multispectral approach to characterize PGC contributions to the ovarian reserve

In zebrafish, absence or an insufficient number of PGCs/germ cells leads to testis development, indicating that there is a threshold number of germ cells needed to establish an ovary (Tzung et al., 2014; Ye et al., 2019). Based on our Germ traCre clonal analysis which revealed that few PGCs give rise to GSCs, we hypothesized that some PGCs give rise to nonrenewing *nos2-/foxl2l-* progenitors to ensure that there are sufficient germ cells to build a biopotential gonad. Moreover, we hypothesized that these progenitors could undergo the first wave of gametogenesis/oogenesis to establish an initial ovarian reserve. Accordingly, oocytes derived from direct progenitors would be nonrenewing and depleted after several matings while oocytes derived later from GSCs would replenish the ovarian reserve and these clones would perdure over reproductive lifespan.

Germ traCre is a highly effective system for analyses of spatially isolated clones within the gonad; however, since all clones are the same color, eggs produced from distinct PGC clones cannot be distinguished. In order to test our hypothesis and parse the contributions of individual PGCs to the ovarian reserve over reproductive lifespan, we developed the multispectral Cre-based transgenic lineage tracing system Germbow, *Tg[ziwi:Zebrabow; cmlc2:mCherry]*. Like Germ traCre, the Germbow construct uses the *ziwi* promoter for stable germline expression of Cre-inducible fluorescent markers and contains an independent transgenesis reporter (*cmcl2:mCherry)* for larval screening (Fig. 4A). To enable multispectral lineage tracing, we used the *Zebrabow* cassette, which encodes dTomato, mCerulean, and EYFP, with flanking floxed stops for Cre-induced random recombination (Pan et al., 2013). The Zebrabow construct was adapted for germline specificity, and also to ensure that any observed colors in embryos were maternally inherited and not due to somatic zygotic expression.

**Fig 4.**
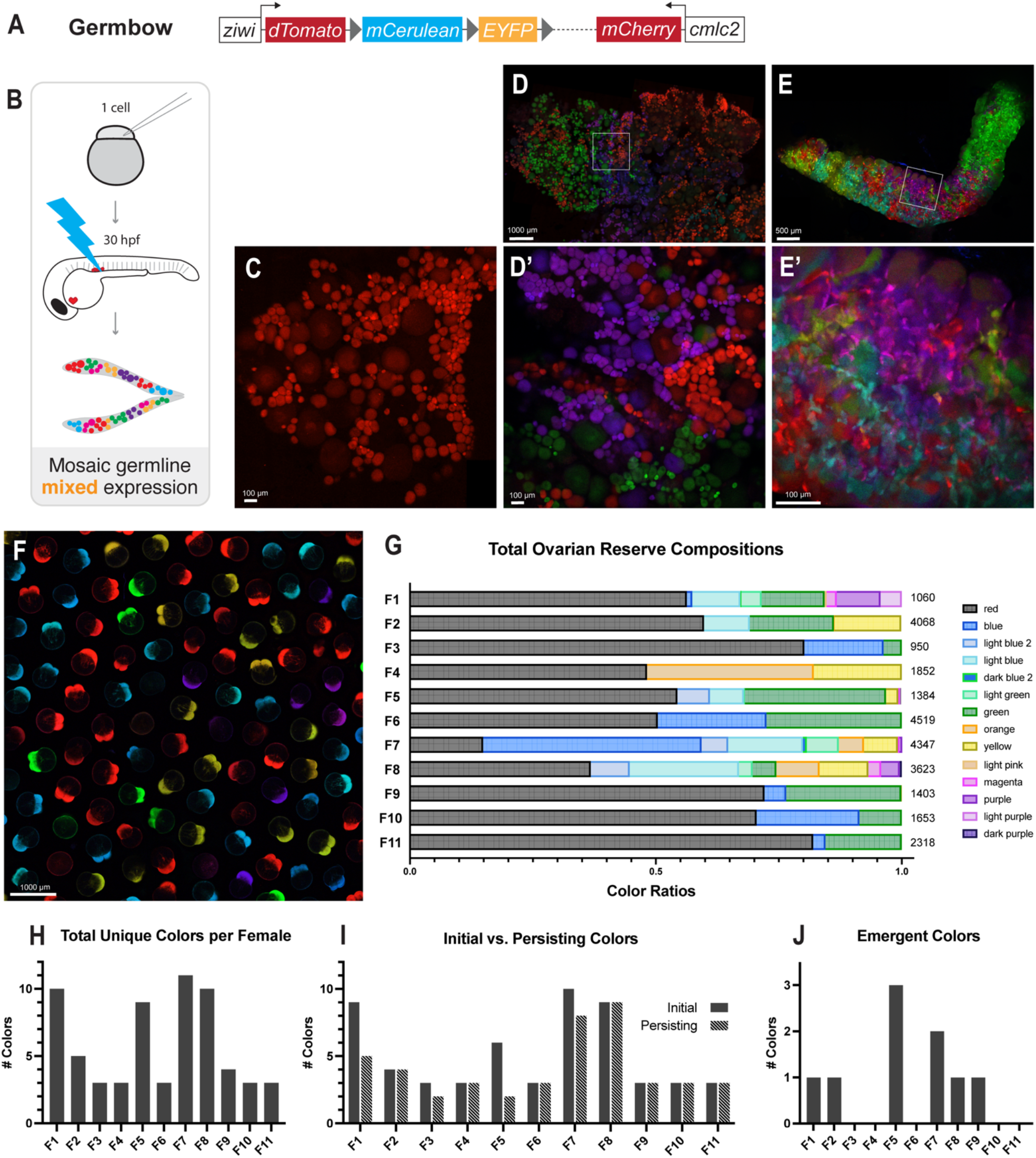
Germbow multispectral germline lineage tracing. **(A)** Schematic of the Germbow transgenic construct. **(B)** Schematic of the Germbow experimental workflow used to generate multispectral mosaic labeling. One-cell stage embryos were injected with *pACre-nos3,3’utr* mRNA, pACre was activated with blue light exposure at 30h to induce random recombination and expression of multispectral markers. **(C)** Control, unactivated ovary expressing dTomato. Scale bar: 100μm. **(D-D’)** Activated adult ovary with mosaic color expression. Scale bars: 1000 μm, 100 μm. **(E-E’)** Activated adult testis with mosaic color expression. Scale bars: 500 μm, 100 μm. **(F)** Representative clutch of eggs/embryos produced from mosaic female matings. Scale bar: 1000 μm. **(G)** Bar graph showing egg color ratios for all mosaic examined Germbow females over the mating assay time course. Total number of eggs analyzed per female indicated. **(H)** Bar graph showing the total number of unique colors observed for each female. **I**. Bar graph comparison of colors initially observed in the first clutch that persisted to the final examined clutch across all females. (**J)** Bar graph of colors that emerged after the first clutch in examined females.

We injected zygotes with a low concentration of *pACre-nos3,3’utr* mRNA and activated dechorionated 30h embryos with blue light to generate mosaic germline labeling (Fig. 4B). Uninjected controls without Cre-induced recombination express only germline dTomato (Fig. 4C). However, in activated gonads, Cre-mediated recombination generates stochastic, combinatorial expression of dTomato, mCerulean, and EYFP (Fig. 4D,E). Notably, these lineage markers were brighter and more stable than mCherry, providing more efficient screening. Since additional IHC/*in situ* labeling is limited in this system, Germbow is not a replacement of Germ traCre. Instead, Germbow augments Germ traCre analyses of early PGC contributions by enabling functional assessment of clonal contributions to the gamete pool over reproductive lifespan. Lines with multiple copies of the transgene yielded numerous color combinations, enabling multilineage clonal analysis of PGC contributions to adult gonads. Interestingly, in contrast with our earlier observations that in the d16 gonad an average of two GSC *nos2*+ clones are initially established (Fig. S1), wholemount of adult ovaries and testes revealed that the majority of examined gonads established more than two clone colors (Figs 4D,E, S3). This suggests that some germ cells in the bipotential gonad adopt GSC fate later in development.

### Individual PGCs can contribute long-term renewing GSCs or nonrenewing progenitors to the adult zebrafish ovarian reserve

To distinguish between PGCs that give rise to renewing GSCs and PGCs hypothesized to produce nonrenewing progenitors, we examined the egg colors of 11 adult female mosaic Germbow fish across several weeks. Females were crossed weekly and freshly-laid clutches of eggs/embryos were imaged using similar parameters to those reported for optimal color resolution in Zebrabow (Pan et al., 2013) (Fig. 4F). Egg colors were used as a proxy for ovarian reserve composition and were quantified over successive matings for each female (Figs 4G, S4). By comparing egg color ratios per clutch over time, we tested our hypothesis that some clones contributed to the ovarian reserve as nonrenewing progenitors which deplete while others contribute to long-term renewing GSCs.

Although the total number of egg colors varied between females (Fig. 4G,H), we observed a shift in egg color ratios for 4/11 females consistent with our hypothesis that some PGCs give rise to nonrenewing progenitors which will deplete over time (Fig. 4I). For example, Female 1 initially produced 9 different egg colors, but only 5 of the initial colors persisted after 7 subsequent matings (Fig. 4I). Notably, the 4 females which displayed reduction in clone colors over time were the only 4 that had more than 5 recombined colors (Fig. 4I,H). This could indicate that color loss was not observed in the other females because recombination efficiency was not high enough to observe loss of single clones. Additionally, multiple clones may have recombined to the same color or indistinguishably similar colors. These limitations have been similarly acknowledged in other papers using the ‘bow’ transgenic lineage tracing approach (Livet et al., 2007; Nguyen et al., 2018; Pan et al., 2013; Weissman and Pan, 2015).

Surprisingly, 5/10 females showed an emergence of a new egg color over time (Fig. 4J), suggesting that some GSCs may be quiescent until the initial ovarian reserve is reduced. For example, magenta eggs emerged in the third clutch from Female 7 and consistently appeared in clutches thereafter (Fig. S4). Female 7 also had the highest recombination efficiency and the greatest number of distinguishable recombination colors, with 11 distinct colors identified across clutches (Fig. 4H).

Combined with our Germ traCre lineage tracing data, our findings indicate that zebrafish PGCs do not all give rise to GSCs as previously thought, but instead differentially contribute to the pre-gametic germline and, ultimately, the adult gonadal reserve (Fig. 5). Although the mechanism(s) driving GSC specification and PGC differentiation remains unclear, individual PGCs directly give rise to multiple germ cell fates.

**Fig 5.**
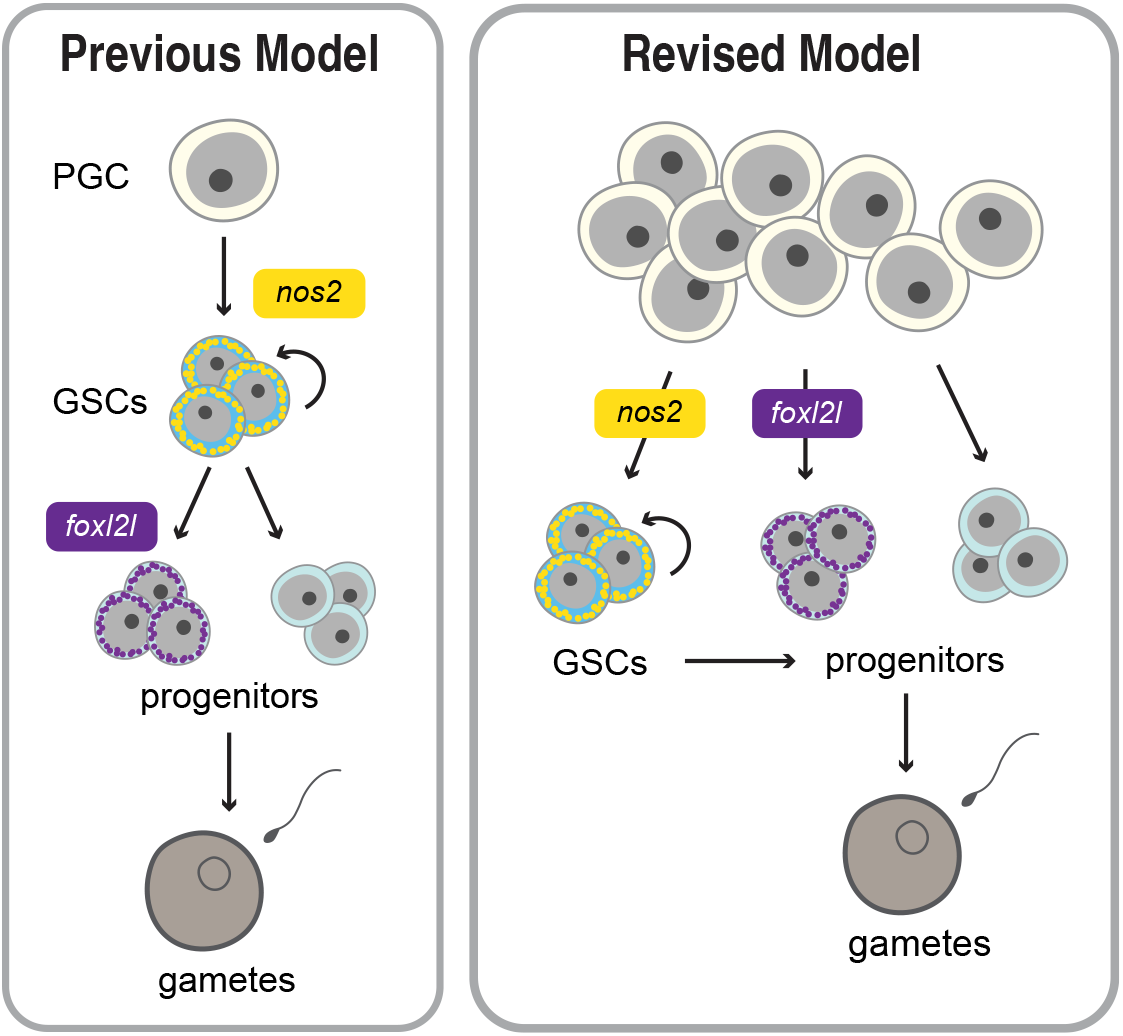
PGC Differentiation Model. (**left**) Schematic depicting the previous model of PGC differentiation and germline development. (**right**) Our revised model of PGC differentiation, where individual PGCs give rise to germline daughter cells with differential expression of the GSC marker *nos2* and the female germline differentiation marker *foxl2l*.

## Discussion

This study establishes two transgenic germline lineage tracing systems in zebrafish, Germ traCre and Germbow, which enable stable, long-term labeling of PGCs and their progeny. Using these systems, we provide functional evidence that individual PGCs directly contribute multiple distinct germ cell fates to form the pre-gametic zebrafish gonad, resulting in differential germline contributions over reproductive lifetime. We demonstrate this hypothesis in the context of the pre-gametic gonad immediately following PGC differentiation, as well as in the adult ovarian reserve. Overall, our findings challenge previous assumptions about zebrafish PGC differentiation and indicate that PGCs contributions are unequal, with some PGCs giving rise to long-term renewing GSCs and others producing nonrenewing progenitors.

Germ traCre lineage tracing of PGCs demonstrated that they can directly give rise to at least four distinct germ cell fates, including the expected and previously known, *nos2*+ GSCs (Cao et al., 2019; Sada et al., 2009), and *foxl2l*+ progenitor populations (Hsu et al., 2024; Kikuchi et al., 2020; Liu et al., 2022; Nishimura et al., 2015; Ren et al., 2024). To our surprise, the largest class of direct PGC descendants identified in our lineage tracing expressed neither *nos2* nor *foxl2l*. Recent scRNAseq of the d26 germline also identified a population of *nos2-/foxl2l-* progenitors (Hsu et al., 2024); however, when we tested the marker identified from this study, *bicc1a*, this did not label *nos2-/foxl2l-* cells. Based on UMAP developmental trajectories of the scRNAseq, *nos2-/foxl2l-* germ cells were assumed to represent an intermediate state between GSCs and *foxl2l*+ ovary-committed progenitors (Hsu et al., 2024). However, our analyses of early clones indicate that *nos2-/foxl2l-* and *foxl2l*+ populations in the pre-gametic gonad are direct descendants of differentiating PGCs rather than an intermediate cell type between *nos2*+ GSCs and *foxl2l*+ progenitors. Separately, Hsu *et al*. (2024, *bioRxiv*) also identified limited double-positive *nos2+/foxl2+* cells in d26 *foxl2l* mutants, but not in WT gonads, leading them to hypothesize that the *nos2+/foxl2+* cells were progenitors reverting to GSC fate in the mutant context. Our data suggests that *nos2+/foxl2+* cells are instead a small, but normal, population in early gonads. Together, our findings indicate PGCs have multiple differentiation paths during germline development, including a direct program from PGC to progenitor that does not involve the previously defined *nos2+* GSCs. Our finding raises questions about what fates are necessary for establishment of the early germline and the roles of these uncharacterized double-negative and double-positive germ cells.

Because *foxl2l* is necessary for ovary fate and dispensable for testis fate (Hsu et al., 2024; Liu et al., 2022; Ren et al., 2024), the *foxl2l*+ clones likely represent ovary-fated progenitors. Given that the d16 gonad is pre-gametic and not yet committed to an ovary or testis fate, one hypothesis is that *nos2-/foxl2l-* germ cells do not express *foxl2l* because they are indeterminant progenitors. Unlike in mammals and birds where sex is determined chromosomally, laboratory zebrafish sex determination is polygenic and influenced by a variety of factors, such as temperature, hormone levels, nutrition, and PGC/germ cell abundance (Abozaid et al., 2025; Dang and Kienzler, 2019; Santos et al., 2017; Slanchev et al., 2005; Tzung et al., 2014; Wilson et al., 2024; Ye et al., 2019). In zebrafish, germ cell abundance is necessary for the establishment of an ovary (Tzung et al., 2014; Ye et al., 2019). Thus, indeterminant progenitors may serve to ensure that there are sufficient germ cells to build a juvenile ovary. Our observation that most PGCs produce transient *nos2-*/*foxl2l-* progenitors may provide an explanation for the seemingly discordant relationship between PGC numbers and early GSC numbers and the dependence of juvenile ovary development on high numbers of PGCs.

Taken together, we propose that ovary formation in zebrafish involves establishment of a primary ovarian reserve akin to the mammalian ovarian reserve, where the first oocytes are produced by transient, nonrenewing progenitors that deplete over time. From our data, it appears that only some PGCs contribute to the primary ovarian reserve. Other PGCs appear to contribute to a secondary renewing reserve arising from GSCs, and this renewable reserve ensures fertility over the zebrafish lifespan. Consistent with this notion, in *nos3* mutant females which lack renewing GSCs, females produce normally patterned embryos for a period of time before becoming sterile males, indicating that zebrafish females can establish a nonrenewing ovarian reserve (Beer et al., 2013; Draper et al., 2007). In light of our lineage tracing analysis, it is conceivable that the initial fertility of *nos3* mutant females is supported by the primary ovarian reserve. The extent to which dysfunctional *nos2+* cells in *nos3* mutant ovaries contribute to the ovarian reserve remains unclear. Taken together, these findings suggest GSCs are not absolutely required for a functional zebrafish ovary but instead enable sustained female fertility over lifespan, something which mammalian ovaries lack.

In contrast to the zebrafish ovary, GSCs appear to be the primary source of germ cells for testis. All known mutants with defects in GSC establishment and/or maintenance develop sterile testes, including *nos2, nos3, ddx4, dazl*, and *piwi* (Beer et al., 2013; Bertho et al., 2021; Cao et al., 2019; Hartung et al., 2014; Leu et al., 2010). Thus, testes development requires GSCs. Our observations that GSCs form along the length of the AP axis of the pre-gametic zebrafish gonad suggests a mechanism for GSC induction that contrasts with the anterior GSC-specifying somatic niche system in invertebrates (Crittenden et al., 2006; Gilboa and Lehmann, 2004; Sheng et al., 2009). This is consistent with wholemount analyses of later developmental stages stained for *nos2*, where GSCs are initially distributed throughout d21 bipotential gonads then become restricted to the germinal ridge in d35 juvenile ovaries (Beer et al., 2013). Together, these observations suggest that, if there are determinative somatic niches in zebrafish, they are likely dynamic or exist throughout the gonad.

Interestingly, although GSCs are initially the minority class in the pre-gametic gonad, with an average of 2 *nos2*+ cells per lobe, our Germbow analyses indicate that adult gonads have more than two renewing populations. This is consistent with the prior quantification of *nos2+* cells at later developmental stages (Beer et al., 2013; Hsu et al., 2024). These observations raise important questions about how GSCs are specified. The emergence of new egg colors over time suggests that a subset of initially *nos2-* cells adopt *nos2* expression and GSC fate over developmental time. Because we found no evidence for fixed inductive niches, we propose that GSC fate is either germ cell-intrinsic or determined by undefined germline and somatic gonad interactions that are not developmentally restricted along the gonadal axes. Based on our finding that PGCs can give rise to different germ cell types without passing through a renewing GSC state, we propose a revised model of early gonad and germline development. Specifically, we propose that differentiation potential varies among PGCs and individual PGCs directly give rise to distinct germ cell types, including renewing GSCs and transient progenitors which are depleted over successive matings. Moreover, these transient progenitors may serve to ensure adequate cells to build a juvenile gonad and provide an early wave of gametes.

The mechanisms underlying zebrafish GSC specification and PGC heterogeneity remain unknown; however, the observed developmental nonequivalence of PGCs may be intrinsic to the PGCs or influenced by interactions with somatic cells. Emerging evidence from both vertebrates and invertebrates indicates that PGC heterogeneity is likely a conserved feature and that PGC fates can be influenced by pathways that precede somatic gonad interactions. In mice, recent studies have indicated that PGC gene expression is heterogeneous and is a potentially competitive mechanism of selection (Jaszczak et al., 2025; Nguyen et al., 2019). Migrating mouse PGCs were found to have transcriptional heterogeneity correlated with their spatial heterogeneity (Jaszczak et al., 2025), which could indicate that non-gonadal somatic interactions drive PGC heterogeneity. Additional support for plasticity of migrating PGCs comes from studies in turtles, wherein PGC commitment to testis or ovary fate is determined by the temperature experienced during the period of PGC migration, prior to interactions with the somatic gonad (Hatkevich et al., 2024; Tezak et al., 2023). Although the mechanisms are not fully understood, PGC fates in some animals appear to be determined prior to arrival and differentiation at the somatic gonad.

Alternatively, since zebrafish PGCs are specified by inheritance of maternal germ plasm, this may be the key contributor to their PGC heterogeneity. In addition to containing factors that specify PGC fate, germ plasm contains mRNAs and RNA-binding proteins, such as *dead end* and Nos3 which drive migration to the gonadal ridge and prevent somatic differentiation (Gross-Thebing et al., 2017; Köprunner et al., 2001; Weidinger et al., 2003). Thus, it is possible that unequal inheritance of maternal germ plasm factors could generate heterogeneity among PGCs (Eno et al., 2016; Hansen et al., 2021; Knaut et al., 2000; Raz, 2002; Rostam et al., 2022; Strasser et al., 2008; Westerich et al., 2023). Close examination in *Drosophila* indicates that germ plasm inheritance is likewise unequally distributed among PGCs (Slaidina and Lehmann, 2017). Moreover, this asymmetry in *Drosophila* PGCs contributes a competitive survival advantage to PGCs with more germ plasm (Slaidina and Lehmann, 2017). If zebrafish PGCs undergo a similar competitive selection, this was not detectable with our lineage tracing approach.

Maternal germ plasm factors could instead also influence transcription or translation of zygotic factors that determine a PGC’s developmental potential. Despite differences in PGC specification between zebrafish and mice (maternal versus zygotic induction), the resulting transcriptional variation and developmental potential of PGCs could be similar (Hackett et al., 2012). Zebrafish PGC transcriptomic profiles have been shown to shift over time (Zhang et al., 2019); however, the extent to which zebrafish PGCs are heterogeneous remains to be determined due to limited availability of single cell sequencing data for this rare cell population. More robust scRNAseq of individual PGC expression profiles over time and functional studies showing their contribution to gonadogenesis and fertility are needed to determine the onset and mechanisms establishing PGC heterogeneity and developmental nonequivalence suggested by our systems.

## Materials and Methods

### Zebrafish strains and handling

All zebrafish (*Danio rerio*) procedures and experimental protocols were performed in accordance with NIH guidelines and were approved by the Icahn School of Medicine at Mount Sinai Institutional Animal Care and Use Committees (ISMMS IACUC; IPROTO202300000039). The following transgenic lines and alleles were used: *Tg[ziwi:stop-mCherry, pGH]*^*mss36*^, and *Tg[ziwi:Zebrabow, pBH]*^*mss69*^.

### Generation of transgenic constructs

Zebrafish transgenesis constructs were generated by Gateway recombination. The Gateway™ LR Clonase™ II Plus Enzyme mix (ThermoFisher, 12538120) was used according to the protocol of the manufacturer to generate all constructs. All vectors have flanking Tol2 transposase sites to enhance transgenesis efficiency (Suster et al., 2009; Thermes et al., 2002). SnapGene software (www.snapgene.com) was used to model and track construct generation. Snapgene files are available upon request. All cloning constructs and materials are listed in **Supplementary Table S1** and all constructs generated in this study are available through Addgene.

#### Germ traCre *(ziwi:stop-mCherry, pGH)*

*The pTol2-ziwi:loxp-stop-loxp-mCherry; cmlc2:gfp* construct (Addgene, #226785) (referred to as Germ traCre and *ziwi:stop-mCherry, pGH*) was generated by recombining the *p5E-ziwi-promoter (gift from Dr. Bruce Draper)* (Leu et al., 2010), *pME-loxp-stop-loxp* (Tol2kit v2.0, #727), and *p3E-mcherry* (Addgene, #108884) (Ariotti et al., 2018) into the *pTol2-R4/R3_cmlc2:gfp* backbone vector (Tol2kit v1.0, #393)(Kwan et al., 2007).

#### Germbow (*ziwi:Zebrabow, pBH*)

The *Zebrabow* cassette from *Zebrabow-GateDest* (Addgene, #118221) (Pan et al., 2013) was excised with Ecl136II (ThermoFisher) and inserted into linearized *pME-MCS_L1/L2* (Tol2kit v2.0) via T4 blunt-end ligation to produce *pME-Zebrabow_L1/L2* (Addgene, #226792). Using Gateway recombination, *pME-Zebrabow_L1/L2* was combined with the *p5E-ziwi-promoter* (Leu et al., 2010) into the *pTol2-R4/R2_cmlc2:mCherry* backbone vector to generate *pTol2-ziwi:Zebrabow; cmlc:mCherry* construct (Addgene #226787) (referred to as Germbow and *ziwi:Zebrabow, pBH*).

#### pACre-nos3,3’utr

The *pCSDest2-pACre-nos3,3’utr* plasmid (Addgene, #226790) used to synthesize *pACre-nos3,3’utr mRNA* was generated by recombining *pCSDest2* (Addgene, #22424) (Villefranc et al., 2007) and *pME-pACre* (Addgene, #226789).

### Generation of transgenic lines

50ng of *Tol2* plasmid DNA for Germ traCre or Germbow was combined with 50ng *Tol2 transposase* mRNA transcribed from *pCS2FA-transposase* (Tol2kit v2.0, #589)(Kwan et al., 2007). One-cell stage embryos were injected with 1-1.5nl of the plasmid/transposase solution. At d4, the selective marker (GFP- or RFP-positive heart) was used to identify potential founders. Embryos expressing transgene markers were raised to adulthood. Founders were identified through crosses with SAT wild type lines. At least 3 lines with stable germline transmission were generated per construct. A subset of embryos from each line were injected with *iCre* mRNA to assess brightness as a proxy for expression levels, and the brightest lines were used for subsequent experiments. Experimental studies of Germ traCre and Germbow were carried out in the representative *ms24* and *ms69* lines, respectively.

### Cre mRNA injections and pACre photoactivation

mRNA was transcribed from *pCSDest2-pACre-nos3,3’utr* and *pME-iCre* (improved Cre; Genbank, KF753697) (Sztal et al., 2015) for use in experimental mosaic labeling and broad activation, respectively. mRNAs were synthesized *in vitro* from SrfI digested plasmids using the mMESSAGE mMACHINE™ SP6 Transcription Kit (Invitrogen, AM1340), according to the protocol of the manufacturer. Immediately following transcription, a portion of the mRNAs were diluted to 250 ng/μL and 1μL aliquots were stored at -80ºC (for 50 ng/μL final concentration in final 5μL injection mix). To induce transgenic marker expression, Germ traCre or Germbow zygotes were injected with 1.5nL mRNA. For pACre photoactivation, injected embryos were dechorionated at 30h, transferred in a single drop to a glass plate, then uniformly exposed to 470 nm blue light for 3 min. under the ZEISS Zoom dissecting scope. No anesthetic was used to allow for embryo movement and exposure of all embryo regions to blue light.

### Hybridization chain reaction (HCR) and immunohistochemistry (IHC)

The HCR-IHC protocol was adapted from the standard Molecular Technologies protocol (Choi et al., 2018). To preserve Germ traCre transgenic mCherry fluorescence signal, samples were fixed in 4% paraformaldehyde (PFA) for 2-4 hours at RT on a shaker, followed by four 5 min. washes with 1x phosphate-buffered saline with 1% Tween-20 (1x PBST). Samples were permeabilized in a 0.25% Proteinase K (ProtK) solution for 5 min. at room temperature (RT) then rinsed twice with 1x PBST, post-fixed in 4% PFA for 5 min., and washed twice for 3 min. each with 1x PBST. Samples were then incubated in 250 μL of pre-warmed probe hybridization buffer (PHB) for 30 min. at 37°C then incubated overnight in probe mixtures, prepared by combining 2-4 μL of each probe into 250 μL of PHB (final concentration 8-16 nM). The following morning, the probe mixtures were removed and stored for reuse and samples were washed four times with warmed PWB for 15 min. at 37°C. Samples were washed twice with 5x saline-sodium citrate with 0.1% Tween-20 (5x SSCT) for 5 min. at RT. Samples were then incubated in Amplification Buffer for 30 min. at RT, followed by overnight static incubation at RT in 250 μL of Amplification Buffer containing the snap-cooled hairpins. On the third day, samples were washed with 5x SSCT at RT twice for 5 min. then four 30 min. washes. In preparation for IHC, samples were washed twice with 1x PBST for 5 min. Samples were then blocked for 2-4 hours at RT in a blocking solution consisting of 5% normal goat serum (NGS), 2% bovine serum albumin (BSA), and 1x PBST. Primary antibody diluted in the blocking solution was then applied overnight at 4°C. On the fourth day, samples were washed six times with 1x PBST at RT for 20 min. each, followed by incubation with secondary antibody diluted in the blocking solution for 2 hours at RT. Finally, samples were washed 4x for 10 min. each with 1x PBST at RT, dissected, and mounted in ProLong™ Diamond Antifade Mountant with DAPI (Invitrogen, P36966) for imaging.

All HCR v3 reagents (probes, hairpins, and buffers) were purchased from Molecular Technologies (Choi et al., 2018). All antibody stainings were carried out as previously described (Leu et al., 2010), with the exception of glycerol clearing. Primary antibodies and their respective dilutions are listed in **Supplementary Table 2**. Secondary antibodies: anti-rabbit/chicken Alexa Fluor dyes AF488, AF568, AF633, AF647, and AF-Cy3 (Molecular Probes) were diluted 1:500. Samples were mounted in ProLong™ Diamond Antifade Mountant with DAPI (Invitrogen, P36962).

Of note, in our experience it was necessary to first complete the HCR FISH protocol before IHC to detect *nos2*, as our initial attempts of first doing IHC then HCR yielded only Vasa and *foxl2l* staining. Limited data was collected for the *foxl2l* single label samples, and the combined data are available in **Supplementary Figure 2**.

### Image acquisition and processing

Sample dissection, mounting, and preliminary imaging was completed using a ZEISS Zoom dissecting scope equipped with Apotome II. Germ traCre and other stained mounted samples were imaged using a Zeiss LSM980 Airyscan II.

Germbow egg/embryo imaging was completed using a Leica SP8 STED 3X. Eggs/embryos were arranged in a monolayer in glass bottom μ-slides (ibidi, 80287). Residual embryo media was carefully pipetted out and embryos were covered with 0.375% low melting point agarose (LMPA; Sigma, A9414) prepared with embryo media. As the agarose set, embryos were gently arranged laterally, with the animal pole laying horizontal to the dish. Slides were left to set on cool benchtop for at least 10 min., until agarose fully solidified. Embryos were imaged using the Leica SP8 STED 3X microscope at 10x magnification fluorescence mode with tiling. The following laser specifications were applied, closely following the Zebrabow protocol (Pan et al., 2013): Argon laser 30%, white light laser 70%.

Image processing was performed in Zen Pro (ZEISS), LAS X (Leica), Fiji/ImageJ, Imaris, and Adobe Illustrator. Germbow egg counting was assisted by Biodock, AI Software Platform (Biodock 2024, available from www.biodock.ai) for identification and categorization of embryos. All results were carefully reviewed and corrected to ensure accuracy.

## Supporting information

Pahima and Marlow Supplemental Data

Pahima and Marlow Supplemental excel file_Germbow eggs

## Acknowledgements

We thank Miranda L. Wilson, Gia Tolins, Paloma Bravo, and past members of the Marlow lab for advice and fruitful discussions throughout this project. We thank our animal care staff at the ISMMS Center for Comparative Medicine for providing excellent animal care of the zebrafish used in this work, and the ISMMS Microscopy CoRE staff for their imaging expertise and assistance with confocal microscopy, supported with funding from NIH Shared Instrumentation Grant (FAIN: S10OD021838). We thank Miranda L. Wilson for kindly providing the *Tol2-ziwi:stop-mcherry; pGH* construct, Dr. Holger Knaut for the rabbit anti-Vasa antibody, and Dr. Bruce Draper for the *nanos3*^*fh49*^ and *Tg[ziwi:GFP]* zebrafish lines. Finally, we are grateful to the Zebrafish Information Network and the Zebrafish International Resource Center for making essential zebrafish resources widely available.

## Author contributions

**M. Pahima**: conceptualization, methodology, investigation, formal analysis and interpretation, writing - original draft, writing - review & editing, visualization, funding acquisition; **F.L. Marlow**: conceptualization, methodology, analysis and interpretation, writing - review & editing, supervision, funding acquisition.

## Data and materials availability

All data and materials will be made freely available upon publication.

