## Supplementary material for "Clonal analysis reveals differential PGC contributions to the early germline and ovarian reserve": Pahima and Marlow Supplemental Data

### Supplementary Materials

**Supplementary Fig. 1: GSC fate location.**

**(A)** Schematic of a d16 gonad (teal, volumetric) with *nos2*+ (yellow) clusters within to represent positioning of clusters. (**B)** Scatter plot showing the number of *nos2*+ cell clusters found per d16 gonad (n=23 gonads examined, n=56 *nos2*+ clusters, mean: 2.45, indicated in red with SD bar). (**C)** Bar graph comparing the positions of *nos2*+ clusters within gonads (periphery n=28, inside n=12, isolated n=16)

**Supplementary Fig. 2: Germ traCre clonal analysis of samples with only *foxl2l* labeling.**

**(A)** Bar graph showing clone type ratios of 195 analyzed clones (*nos2*+: 17, 8.72%; *foxl2l*+: 63, 32.31%; *nos2*+/*foxl2l*+: 4, 2.05%; *nos2*-/*foxl2l*-: 50, 25.64%; *foxl2l*-: 61, 31.28%). **(B)** Bar graph showing the distribution ratios of clone types along the gonadal AP axis. To quantify this, gonads separated into 4 quadrants (Q1-Q4) for analysis. Total number of clones per quadrant labeled. **(C)** Scatter plot of clone size distributions of 217 clones along the gonadal AP axis. Means indicated in red with black SD bars (Q1: 46, 4.85 ± 4.93; Q2: 46, 5.00 ± 6.18; Q3: 78, 3.78 ± 6.54; Q4: 67, 4.27 ± 6.39). **(D)** Scatter bar plot of clone sizes between clone types, SD bars indicated (*nos2*+: 17, 2.18 ± 1.19; *foxl2l*+: 28, 3.68 ± 3.32; *nos2*+/*foxl2l*+: 4, 2.50 ± 0.58; *nos2*-/*foxl2l*-: 50, 1.92 ± 1.58; *foxl2l*-: 61, 4.51 ± 6.57). **(E)** Bar plot showing ratio of one-cell clones to multi-cell clones per clone type (*nos2*+: 7:10, 41.18%; *foxl2l*+: 15:48, 23.81%; *nos2*+/*foxl2l*+: 0:4, 0%; *nos2*-/*foxl2l*-: 32:18, 64%; *foxl2l*-: 24:37, 39.34%).

**Supplementary Figure 3: Number of colors observed in Germbow adult mosaic testes.**

**(A)** Germbow mosaic testes. Scale bar: 500 µm. **(B)** Scatter plot showing the number of unique colors identified across wholemount mosaic Germbow testes (n=6). Mean indicated in red with black SD bars (4.33 ± 2.73).

**Supplementary Figure 4: Germbow egg color lineage tracing bar graph time course representations.**

Bar graphs representing egg color ratios over mating assay time course for **(A)** females with Cre low recombination and **(B-E)** females with high recombination.

**Supplementary Table 1: Cloning reagents and sources for the Germ traCre, Germbow, and pACre-nos3 constructs.**

**Supplementary Table 2: Antibodies and probes for fluorescent staining.**

**Supplementary Data: Spreadsheet of mosaic Germbow female egg data.**

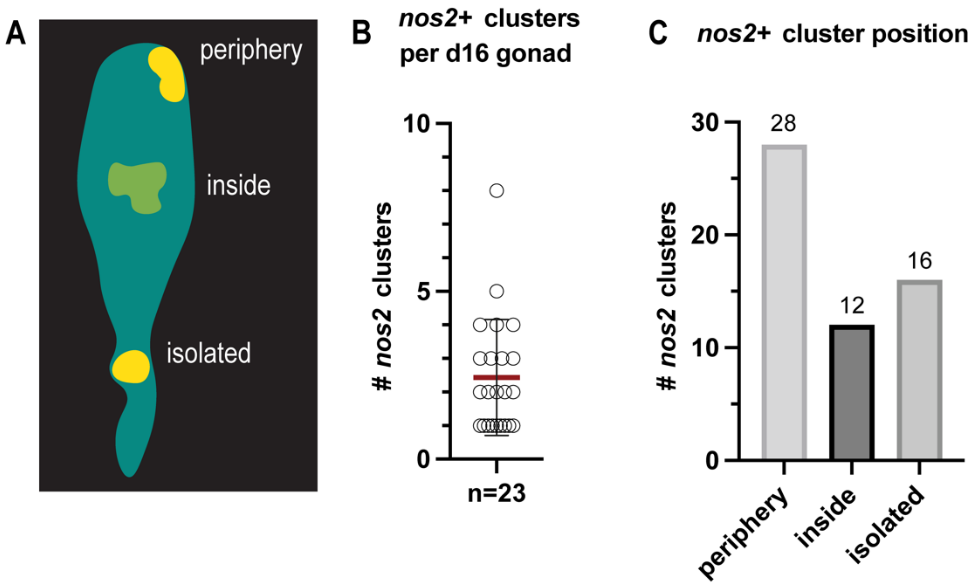

**Supplementary Fig. 1: GSC fate location. (A)** Schematic of a d16 gonad (teal, volumetric) with *nos2*+ (yellow) clusters within to represent positioning of clusters. (**B)** Scatter plot showing the number of *nos2*+ cell clusters found per d16 gonad (n=23 gonads examined, n=56 *nos2*+ clusters, mean: 2.45, indicated in red with SD bar). (**C)** Bar graph comparing the positions of *nos2*+ clusters within gonads (periphery n=28, inside n=12, isolated n=16)

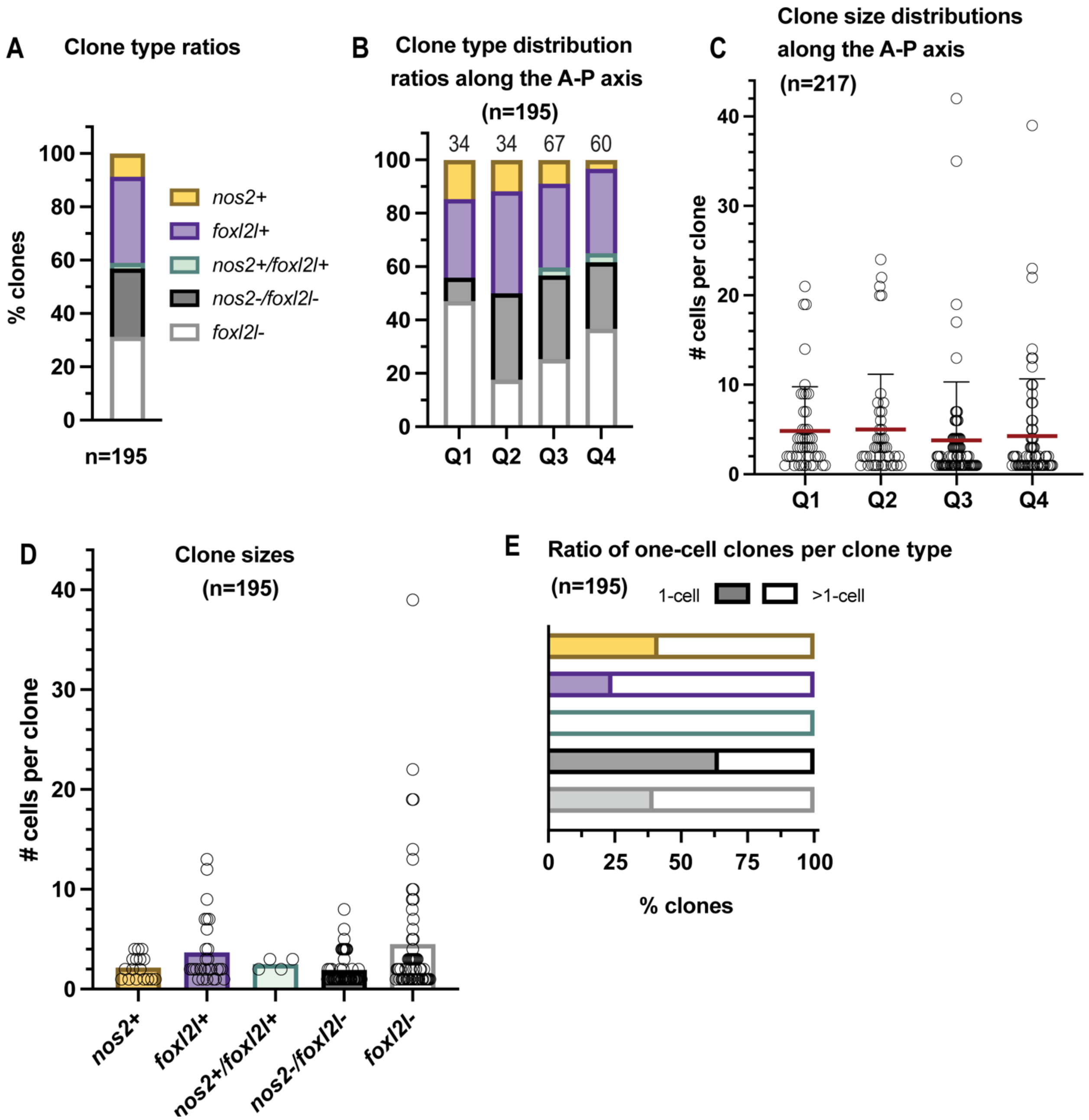

**Supplementary Fig. 2: Germ traCre clonal analysis of samples with only *foxl2l* labeling. (A)** Bar graph showing clone type ratios of 195 analyzed clones (*nos2*+: 17, 8.72%; *foxl2l*+: 63, 32.31%; *nos2*+/*foxl2l*+: 4, 2.05%; *nos2*-/*foxl2l*-: 50, 25.64%; *foxl2l*-: 61, 31.28%). **(B)** Bar graph showing the distribution ratios of clone types along the gonadal AP axis. To quantify this, gonads separated into 4 quadrants (Q1-Q4) for analysis. Total number of clones per quadrant labeled. **(C)** Scatter plot of clone size distributions of 217 clones along the gonadal AP axis. Means indicated in red with black SD bars (Q1: 46, 4.85 ± 4.93; Q2: 46, 5.00 ± 6.18; Q3: 78, 3.78 ± 6.54; Q4: 67, 4.27 ± 6.39). **(D)** Scatter bar plot of clone sizes between clone types, SD bars indicated (*nos2*+: 17, 2.18 ± 1.19; *foxl2l*+: 28, 3.68 ± 3.32; *nos2*+/*foxl2l*+: 4, 2.50 ± 0.58; *nos2*-/*foxl2l*-: 50, 1.92 ± 1.58; *foxl2l*-: 61, 4.51 ± 6.57). **(E)** Bar plot showing ratio of one-cell clones to multi-cell clones per clone type (*nos2*+: 7:10, 41.18%; *foxl2l*+: 15:48, 23.81%; *nos2*+/*foxl2l*+: 0:4, 0%; *nos2*-/*foxl2l*-: 32:18, 64%; *foxl2l*-: 24:37, 39.34%).

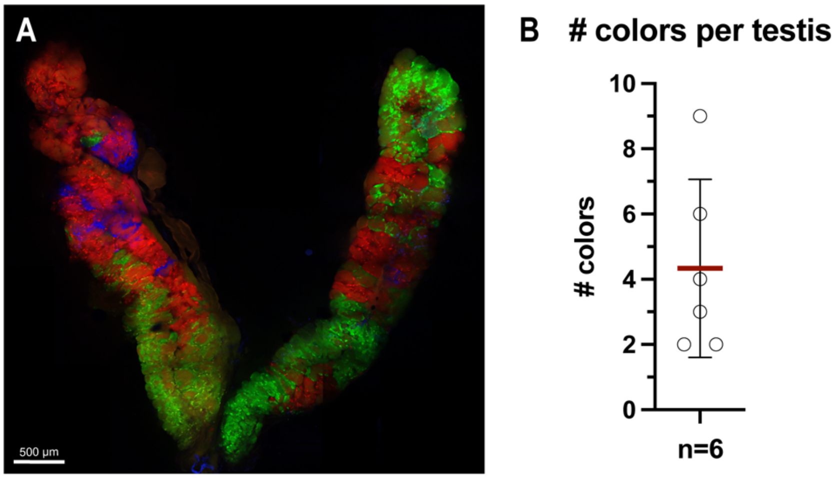

**Supplementary Figure 3: Number of colors observed in Germbow adult mosaic testes. (A)** Germbow mosaic testes. Scale bar: 500 µm. **(B)** Scatter plot showing the number of unique colors identified across wholemount mosaic Germbow testes (n=6). Mean indicated in red with black SD bars (4.33 ± 2.73).

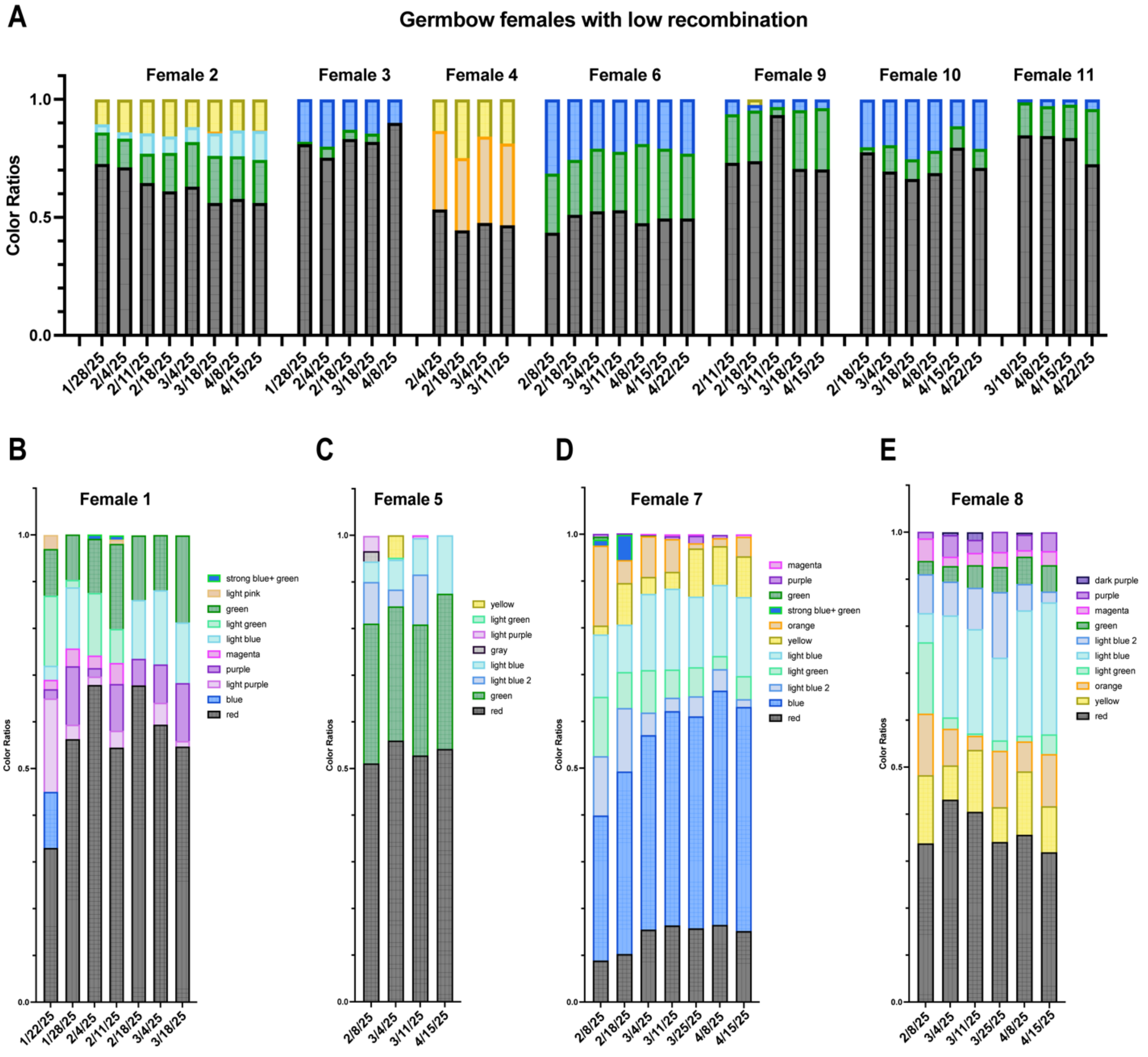

**Supplementary Figure 4: Germbow egg color lineage tracing bar graph time course representations.** Bar graphs representing egg color ratios over mating assay time course for **(A)** females with Cre low recombination and **(B-E)** females with high recombination.

**Supplementary Table 1**

| Plasmids +A2:E24used | Plasmids used | Plasmids used | Plasmids used | Plasmids used |
|---|---|---|---|---|
| *pCS2FA-transposase* | Tol2kit v1.0 | #396 | (Kwan *et al.* , 2007) | |
| *pME-iCre (improved Cre)* | Tol2kit v2.0 | #589 | (Sztal *et al.* , 2015) | |
| ***pTol2-ziwi:loxp-stop-loxp-mcherry; cmlc2:gfp*** | generated in this study, Addgene | #226785 | | *ziwi:stop-mCherry, pGH* (**Germ traCre**) |
| *pTol2-R4/R3_cmlc2:gfp* | Tol2kit v1.0 | #393 | (Kwan *et al.* , 2007) | *pDestTol2CG* |
| *p5E-ziwi* | gift from Dr. Bruce Draper | | (Leu and Draper, 2010) | |
| *pME-loxp-stop-loxp* | Tol2kit v2.0 | #727 | | |
| *p3E-mcherry* | Addgene, Dr. Rob Parton | #108884 | (Ariotti *et al.* , 2018) | |
| ***pCSDest2-pACre-nos3,3'utr*** | generated in this study, Addgene | #226790 | | *p+E12BH-R4/R2+A2:E24* |
| *pCSDest2* | Addgene, Dr. Nathan Lawson | #22424 | (Villefranc et al., 2007) | |
| *p3E-nos3,3'utr* | Gift from Dr. Erez Raz | | (Blaser et al., 2005) | |
| *pME-pACre* | generated in this study, Addgene | #226789 | | |
| *pME-MCS (Multi Cloning Site)* | Tol2kit v2.0 | | | |
| *pcDNA3.1-pACre* | Addgene, Dr. Moritoshi Sato | #122960 | (Kawano et al., 2016) | |
| ***pTol2-ziwi:Zebrabow; cmlc2:mCherry*** | generated in this study, Addgene | #226787 | | *ziwi:Zebrabow, pBH* (**Germbow**) |
| *pTol2Dest-R4/R2_cmlc2:mCherry* | Marlow lab | | (Heim et al., 2014) | *pBH-R4/R2* |
| *p5E-ziwi* | gift from Dr. Bruce Draper | | (Leu and Draper, 2010) | |
| *pME-Zebrabow* | generated in this study, Addgene | #226792 | | |
| *Zebrabow_GateDest* | Addgene, Dr. Albert Pan | #118221 | (Pan et al., 2013) | |
| *pME-MCS (Multi Cloning Site)* | Tol2kit v2.0 | | | |

**Supplementary Table 2**

| Primary antibodies and probes | Dilution/Conc. | Source |
|---|---|---|
| Rabbit anti-Ddx4/Vasa (polyclonal) | 1:2000 | PMID:10811828 |
| Chicken anti-Ddx4/Vasa (polyclonal) | 1:2000 | PMID: 30653507 |
| *nanos2* | 16nM | Molecular Instruments (PRB905) |
| *foxl2l* | 8nM | Molecular Instruments (PRD331) |
| *rec8a* | 16nM | Molecular Instruments (PRE326) |
| *bicc1a* | 16nM | Molecular Instruments (RTP763) |
